# Multimodal Brain Signal Complexity Predicts Human Intelligence

**DOI:** 10.1101/2022.06.25.497602

**Authors:** Jonas A. Thiele, Aylin Richter, Kirsten Hilger

## Abstract

Spontaneous brain activity builds the foundation for human cognitive processing during external demands. Neuroimaging studies based on functional magnetic resonance imaging (fMRI) identified specific characteristics of spontaneous (intrinsic) brain dynamics to be associated with individual differences in general cognitive ability, i.e., intelligence. However, fMRI research is inherently limited by low temporal resolution, thus, preventing conclusions about neural fluctuations within the range of milliseconds. Here, we used resting-state electroencephalographical (EEG) recordings from 144 healthy adults to test whether individual differences in intelligence (Raven’s Advanced Progressive Matrices scores) can be predicted from the complexity of temporally highly resolved intrinsic brain signals. We compared different operationalizations of brain signal complexity (multiscale entropy, Shannon entropy, Fuzzy entropy, and specific characteristics of microstates) regarding their relation to intelligence. The results indicate that associations between brain signal complexity measures and intelligence are of small effect sizes (*r* ~ .20) and vary across different spatial and temporal scales. Specifically, higher intelligence scores were associated with lower complexity in local aspects of neural processing, and less activity in task-negative brain regions belonging to the defaultmode network. Finally, we combined multiple measures of brain signal complexity to show that individual intelligence scores can be significantly predicted with a multimodal model within the sample (10-fold cross-validation) as well as in an independent sample (external replication, *N* = 57). In sum, our results highlight the temporal and spatial dependency of associations between intelligence and intrinsic brain dynamics, proposing multimodal approaches as promising means for future neuroscientific research on complex human traits.

**Significance Statement:** Spontaneous brain activity builds the foundation for intelligent processing - the ability of humans to adapt to various cognitive demands. Using resting-state EEG, we extracted multiple aspects of temporally highly resolved intrinsic brain dynamics to investigate their relationship with individual differences in intelligence. Single associations were of small effect sizes and varied critically across spatial and temporal scales. However, combining multiple measures in a multimodal cross-validated prediction model, allows to significantly predict individual intelligence scores in unseen participants. Our study adds to a growing body of research suggesting that observable associations between complex human traits and neural parameters might be rather small and proposes multimodal prediction approaches as promising tool to derive robust brain-behavior relations despite limited sample sizes.

## Introduction

Intelligence is one of the most important predictors of crucial life outcomes, such as academic achievement, health, and longevity (Calvin et al., 2011; Deary et al., 2004; Sternberg, 1997). Identifying biomarkers of individual differences in intelligence in brain function and brain structure is an ongoing goal of neuroscientific research (Basten et al., 2015; Haier, 2017; Jung and Haier, 2007). Spontaneous brain activity observed during the so-called resting state, i.e., in the absence of an instructed task, is suggested to reflect intrinsic neural communication. Individual differences in such intrinsic neural dynamics are assumed to possess trait character (Hilger and Markett, 2021) and to determine major parts of individual neural processing during external demands (Cole et al., 2014; Schultz and Cole, 2016; Thiele et al., 2022). Different characteristics of intrinsic brain activity have also been identified as biomarkers of a person’s general cognitive ability level, mostly operationalized as intelligence (Hilger and Sporns, 2021).

The majority of this research is, however, based on neuroimaging techniques (functional magnetic-resonance imaging) with limited temporal resolution (~1Hz). Electroencephalography (EEG) allows to study intrinsic neural dynamics with much higher temporal resolution (~250Hz) and measures of intrinsic brain dynamics that have been related to individual differences in intelligence include graph measures (Langer et al., 2012), coherence (Cheung et al., 2014), and theta-gamma cross-frequency coupling (Pahor and Jaušovec, 2014).

Besides those conventional EEG measures, the complexity (or unpredictability) of spontaneous brain dynamics contains additional information about intrinsic predispositions for information processing across functional brain networks (McDonough and Nashiro, 2014). Sufficient complexity of neural signals was hypothesized to constitute the basis for efficient neural communication (de Pasquale et al., 2016) and higher complexity was related to increased information processing capacity (Heisz and McIntosh, 2013). Further, individual variations in brain signal complexity have been associated with differences in creativity (Kaur et al., 2021), the genetic risk for psychological diseases (Li et al., 2008), and also with differences in intelligence (Dreszer et al., 2020; Lutzenberger et al., 1992). However, across-study findings on relations between individual differences in intelligence and intrinsic brain signal complexity are heterogeneous. Higher intelligence has been associated with higher complexity (Jaušovec and Jaušovec, 2003; Stankova and Myshkin, 2016; Thatcher et al., 2005), Dreszer et al. (2020) found both positive and negative associations, and other studies could not find any significant relation (Anokhin et al., 1999; Ueno et al., 2015). Sample characteristics (Dreszer et al., 2020), varying complexity measures (Ferenets et al., 2006; Goldberger et al., 2002), and differences in the considered type of neural complexity, i.e., complexity that has been linked to local (short-range) vs. global (long-range) neural processing (Courtiol et al., 2016; Dreszer et al., 2020; Vakorin et al., 2011) were proposed as contributing to this heterogeneity. Consequently, results of the existing studies are difficult to compare and highlight the need for a holistic framework investigating the relation between intelligence and brain signal complexity with various measures, on multiple spatial and temporal scales and in different study samples.

Most studies investigated brain signal complexity exclusively using different entropy measures. Properties of EEG microstates, defined as spontaneous spatial brain activity patterns (Lehmann et al., 1987) provide complementary insights, i.e., into the spatial dimension of neural dynamics’ complexity. Notably, appearance, stability, and variability of microstates have been associated with variations in intelligence (Liu et al., 2020; Santarnecchi et al., 2017). However, entropy and microstate measures have only been examined in separation of each other but not in a combined analysis approach, so far.

In this study we used resting-state EEG of 144 adults to investigate associations between individual differences in intelligence and variations in brain signal complexity. We developed a framework that combines multiple measures of entropy and microstates, multiple brain regions, and multiple temporal scales to show that individual intelligence scores can significantly be predicted by a multimodal brain complexity model. Significant prediction was achieved within the sample via internal cross-validation as well as in an external replication sample.

## Materials and Methods

### Preregistration

Before data analysis, we preregistered our correlative analyses, sample size, and variables of interest in the Open Science Framework. The preregistration can be accessed under https://osf.io/9sp36. Please note that the predictive analyses (see below) were not preregistered, as they were developed afterwards to overcome the limitations inherent to the low effects sizes of single associations observed in the preregistered analyses.

### Participants

Main analyses were conducted on data from 150 healthy, adult, and right-handed males. The size of the sample was determined by a combination of a priori power calculations and monetary feasibility. Specifically, the minimal sample size to detect effect sizes of *r* = .25 (for meta analysis see Nuijten et al., 2020) with a power of 80% (*α* = .05) was 123, setting the lower limit for our study; our monetary resources allowed us to recruit 27 additional participants. The decision to stop at 150 participants was drawn before data acquisition started. Recruitment was performed through the online study registration system of the University of Würzburg as well as via flyers and various bulletin board websites. Study participation was monetarily compensated with 10€/h. Students with a Major or Minor study subject in Psychology were excluded. All participants had self-reported normal or corrected-to-normal visual acuity, no use of drug substances, no history of chronic pain, no psychiatric or neurological diseases, and normal cardiovascular and endocrinological conditions. Note, that the reason for the exclusively male sample and the above-mentioned exclusion criteria was that data acquisition took place as part of a larger project, in which, beyond others, differences in stress hormone concentrations were analyzed and differences in menstruation cycles needed to be prevented.

All study procedures were approved by the local ethic committee (Psychological Institute, University of Würzburg, Germany, GZEK 2020 −18), and informed consent in accordance with the declaration of Helsinki was obtained from all participants. After the exclusion of six participants due to excessive artefacts (see below), the final sample consisted of 144 subjects (18-35 years, mean age = 24.9 years).

### Intelligence

Intelligence was assessed in group settings (up to five participants) with Raven’s Advanced Progressive Matrices (RAPM, Raven and Court, 1998) under timed conditions (time limit of 40 minutes). RAPM scores ranged between 14 and 36 (*M* = 26.90, *SD* = 4.48) corresponding to intelligence quotient (IQ) scores between 65 and 139 (*M* = 100.49, *SD* = 15.41). RAPM sum scores were used in all analyses as the variable of interest (see Fig. 1A for RAPM sum score distributions).

**Figure 1.**
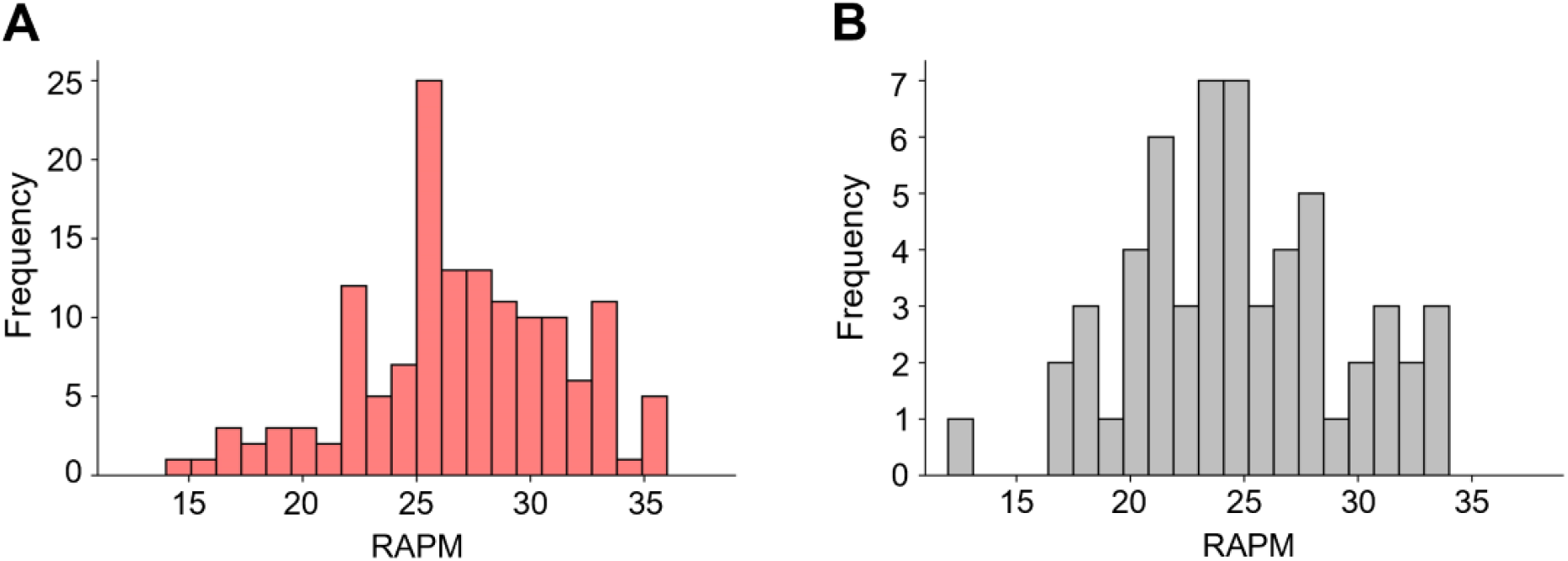
Frequency distributions of intelligence scores. Frequencies of individual intelligence scores (RAPM sum scores, Raven and Court, 1998) are depicted for (**A**) the main sample and (**B**) the replication sample. *Figure Contributions:* JT and KH designed research; JT, AR, KH performed research; JT & AR analyzed data.

### Electroencephalographical Recordings and Preprocessing

Electroencephalography (EEG) data were recorded during five minutes of eyes-closed resting state. Participants sitting on a chair in a sound-shielded room, were instructed to stay relaxed, to prevent motion, to not engage in specific thoughts during the measurement period, to close their eyes, and to remain awake. The EEG was monitored by the experimenter throughout the recording to detect irregularities. Due to the Covid-19 pandemic, medical masks had to be worn during the lab visit. Data were recorded with Brainvision Recorder using 28 active Ag/AgCl electrodes (arranged in a 10–20 layout) and an actiChamp amplifier (Brain Products GmbH, Gilching, Germany). FCz served as online reference, and AFz as ground. The sampling rate was 1000 Hz, impedance levels were kept below 5 kOm, and a low pass filter of 250 Hz was applied during acquisition (notch filter on). Two additional electrodes were placed below the left (SO1) and the right (SO2) eye to record ocular artifacts, and mastoid electrodes were placed behind both ears (M1, M2). Preprocessing of EEG data were conducted in Python using Python MNE (Gramfort et al., 2013) and Pyprep (Bigdely-Shamlo et al., 2015).

For preprocessing, EEG data was bandpass filtered (0.1 – 40 Hz) and down sampled to 250 Hz. EEG channels with NaN (not a number) values, with flat signal, with abnormally high or low overall amplitudes detected by an absolute z-score 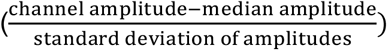 larger than 5, and channels whose time signal correlated too low with signals of the other channels within time segments of one second length (maximum Pearson correlation between the signal of one channel in a given time segment and the signal of another channel within the same segment *r* < .4 in more than 1% of all segments) were interpolated (Pyprep, Bigdely-Shamlo et al., 2015). An average reference was estimated with signals from all 28 scalp electrodes and all signals were re-referenced to this average reference. Subsequently, eye movements were corrected via independent component analysis (ICA) using MNE’s ICA preprocessing tool FastICA (20 components). Eye movement and muscle artefact components were manually identified on ten exemplary subjects through visual inspection and through matching with typical EOG activity component maps (Jung et al., 2000). For all other participants, components strongly correlated with these artefact components were identified (using MNE’s corrmap tool, Campos Viola et al., 2009) and removed. The first and last 10 seconds of the signal were discarded. Finally, data were segmented into 2s epochs and epochs exceeding 1e-4 V were removed. Participants with more than one third of the epochs removed were excluded from further analysis (six participants).

### Intrinsic Brain Signal Complexity

Three entropy measures were calculated to gain a holistic picture of intrinsic brain signal complexity (i.e., uncertainty of the EEG signal, see Figure 2 for a simplified overview):

1. Shannon entropy (Shannon, 1948) measures the uncertainty of one event *x_i_* (single voltage amplitude of the EEG signal in the time domain) based on the probability *P* distribution of all events *x* (all voltage amplitudes of the EEG signal in the time domain) of the EEG signal:

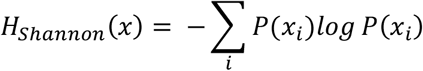
2. Fuzzy entropy (Azami et al., 2019; De Luca and Termini, 1972) depicts the quantity of information expressed by the EEG signal based on the probability *P* that two data point patterns (patterns of consecutive voltage amplitudes of the EEG signal in the time domain) *x_i_* and *x_j_* of the length *m* continue to be similar after adding a further datapoint. First, consecutive datapoints with length *m* (or *m* + 1) are extracted from the signal (signal length = *N*) as template vectors *x* with a time delay *d* (here *d* = 1) between the vectors. Then, a baseline is removed from all vectors *x* and distances Δ (Chebyshev distance) between all vectors of length *l* = *m* (or *l* = *m + 1*) are calculated. The degree of similarity *θ_ij_* between each of two patterns *x_i_* and *x_j_* of lengths *l* = *m* (or *l* = *m* + 1) is determined in respect to the tolerance *r* with a fuzzy membership function 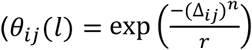, here *n* = 1, and *r* = 0.2). Next, based on these degrees of similarity, the total probabilities that two patterns of lengths *l* = *m* (or *l* = *m* + 1) match are calculated:

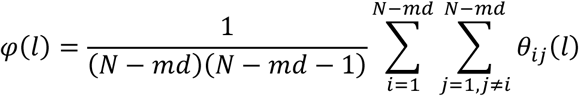 Fuzzy entropy is then determined in respect to the functions *φ(m)* and *φ*(*m* + 1):

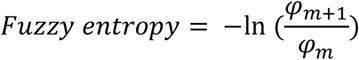
3. Multiscale entropy (Costa et al., 2005, 2002) is an extension of sample entropy (Richman and Moorman, 2000) and goes beyond traditional entropy measures in capturing the regularity (predictability) of time series on multiple time scales. First, multiple coarse-grained time series of the EEG signal (voltage amplitudes in the time domain) are constructed by averaging amplitudes of consecutive data points within non-overlapping time segments of length *τ* (with *τ* representing the time scale). Then, sample entropy is determined for each coarse-grained signal (scale). Sample entropy is computed similar to Fuzzy entropy (see above), with the differences that no baseline correction of the template vectors is performed before calculating the Chebyshev distances between them and that the similarity between two template vectors of lengths *l* = *m* (or *l* = *m* + 1) is determined in respect to the tolerance *r* (here *r* = 0.2 times the standard deviation of the signal) by:

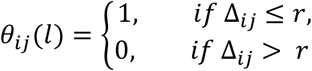 For mathematical insights and more detailed descriptions see Richman and Moorman (2000), and Valencia et al. (2019).

**Figure 2.**
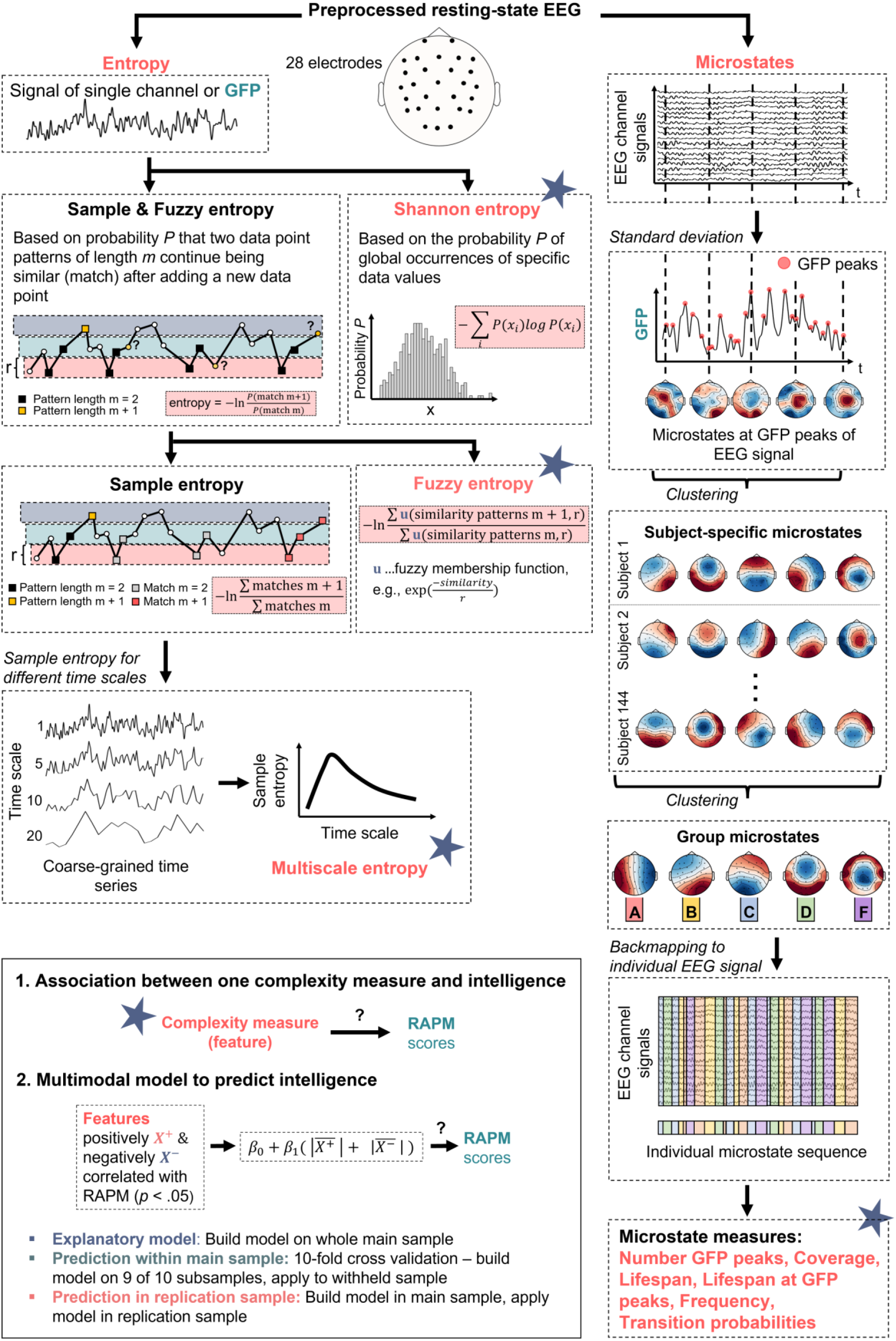
Schematic overview of the analyses to investigate relations between brain signal complexity and intelligence (RAPM sum scores; Raven and Court, 1998). First, from the preprocessed resting-state electroencephalography (EEG) signals of 144 participants different entropy measurers (Shannon entropy, Fuzzy entropy, multiscale entropy, see left branch) and microstate measures (number of GFP peaks, coverage, lifespan, lifespan at GFP peaks, frequency, and transition probabilities between microstates, see right branch) were computed. Entropy measures were determined for the GFP of the EEG signal and for each EEG channel, respectively; for multiscale entropy, sample entropy was calculated for different coarse-grained time series (scales). For microstate analyses, individual brain signals at GFP peaks were clustered (k-means) over time into five individual spatial mean maps (microstates). The individual mean maps of all participants were then clustered into five group microstates. These group-average microstates were then mapped back onto the individual EEG time series resulting in a sequence of group microstates for each individual from which different microstate measures were derived. As depicted in the lower left box, first, all associations between intelligence and single complexity measures were determined. Second, a multimodal model was constructed from measures of brain signal complexity significantly positively or negatively correlated (*p* < .05, uncorrected) with intelligence. It was tested a) to which extent the model explains intelligence scores in the main sample, b) with which accuracy the model predicts intelligence scores within the main sample (10-fold cross-validation), and c) with which accuracy the model predicts intelligence scores in the replication sample (out-of-sample prediction, model build in the main and applied to the replication sample). GFP, Global field power. Note that all equations are simplified for illustration purposes, see Methods for more details. *Figure Contributions:* JT and KH designed research; JT created the figure.

All entropies measures were computed on the global field power as well as on each of the 28 EEG channels separately. MSE was calculated for the time scales *τ*∈ [1..20] (Costa et al., 2005), which, for the used sample rate of 250 Hz, corresponds to time segments with a size of 4 to 80 milliseconds.

For analyses of EEG microstates (Lehmann et al., 1987; Michel and Koenig, 2018) first, the global field power (GFP, Lehmann, 1971) of each subject-specific EEG was calculated as the standard deviation of the average-referenced signal across all electrodes. Second, individuals’ activity patterns of the 28 scalp electrodes occurring at the peaks of the GFP-signal were clustered (via modified k-means, Pascual-Marqui et al. 1995) into five spatial mean maps (individual microstates, see Michel and Koenig, 2018) with the criteria to maximize the global variance of the subject-specific EEG-signal that can be explained by these maps (GEV, 1000 iterations, segmentation with maximal GEV was chosen, Poulsen et al., 2018). Third, all subjects’ mean maps were clustered with modified k-means clustering into five group maps (group microstates), again with the criteria to maximize the total explained variance. Fourth, the derived group microstates were backfitted to each individuals’ original EEG time-series. Specifically, each time point was assigned to the group microstate that expresses the highest correlation with the individual-specific activity map at this specific time point (Liu et al., 2020; Santarnecchi et al., 2017). This step resulted in a subject-specific sequence of group microstates that presents the input for the calculation of six different individual-specific measures (Brodbeck et al., 2012; e.g., Koenig et al., 2002; Seitzman et al., 2017): 1. Coverage (proportion of total number of time points assigned to a specific group microstate), 2. Lifespan (mean duration of a microstate, i.e., mean number of consecutive time points assigned to the same group microstate), 3. Frequency (how often a microstate appears, i.e., count of instances a group microstate appears for the first time, after time points assigned to a different group microstate), 4. Lifespan at GFP peaks (mean duration of a microstate, i.e., mean number of consecutive GFP peaks assigned to the same group microstate), 5. Number of GFP peaks, and 6.Transition probabilities (likelihood of a group microstate to continue or to transition into another group microstate).

Entropy and microstate measures together resulted in 709 brain signal complexity metrics per participant, i.e., global Shannon entropy, global Fuzzy entropy, and global sample entropy at 20 scales (22 global entropy measures calculated at the GFP of the EEG signal); 2 × 28 channel-wise Fuzzy entropy and Shannon entropy; 28 × 20 channel- and scale-wise sample entropy; number of GFP peaks; coverage, life span, frequency, and lifespan at GFP peaks for five microstates (4 × 5 measures); 5 × 5 transition probabilities between microstates; 5 × 5 transition probabilities between microstates at GFP peaks. For a simplified and schematic overview of the measures and analyses see Figure 2. To investigate the covariance pattern of the different neural complexity measures, an exploratory factor analyses with oblique rotation (promax) and minimal residual (MINRES) fitting was performed. The number of factors was determined by parallel analysis (Horn, 1965).

### Correlative Associations Between Intrinsic Brain Signal Complexity and Intelligence

Relationships between complexity measures and intelligence were assessed with partial Pearson correlations by controlling for age (Pearson correlation between intelligence and age: *r = .04, p =.657*) and the number of epochs removed due to artefacts (Pearson correlation between intelligence and number of removed epochs: *r = .14, p = .083).* Statistical significance was accepted as *p* < .05. P-values were corrected via false discovery rate (FDR) or permutation testing. Specifically, significances of relations between intelligence and a) channel-wise entropy were FDR corrected for the number of channels, i.e., 28; b) microstate coverage, frequency, and lifespan were FDR corrected for number of measures and number of microstates, i.e., 4 × 5 = 20; c) microstate transition probabilities were FDR corrected for the number of possible transitions, i.e., 5 × 5 = 25 respectively; and d) MSE were assessed via permutation testing (minimal cluster size for *p* < .05 was assessed by computing the probability of the occurrence of each cluster size across 100 permutations). Note that due to the explorative character of this study (high number of variables explored) and for hypotheses-generating purposes, in addition to corrected *p*-values, indicated as *p(adj)*, effects without correction for multiple comparisons are also reported, indicated as *p*.

### Predicting Intelligence from Multimodal Brain Signal Complexity

To predict intelligence from brain signal complexity, we developed a multimodal approach that combines multiple measures (entropies and microstate measures) into a single multivariate model. In analogy to the most popular prediction frameworks in neuroimaging (connectome-based predictive modeling, CPM, Finn et al., 2015; Shen et al., 2017), the input features of this model were the means of z-normalized complexity measures positively X^+^ and negatively X^-^ correlated with intelligence (*p* < .05):

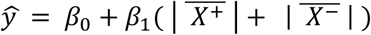

with *β*_0_ = 0 and *β*_1_ = 1 and the predicted intelligence score *ŷ*. To account for multicollinearity and the high proportion of MSE measures, MSE measures were averaged within spatial and temporal clusters (Dreszer et al., 2020). Specifically, we combined electrodes into eight spatial clusters: frontopolar (Fp1, Fp2), frontocentral (FC1, FC2, Fz, FC5, FC6), frontal (F3, F4, F7, F8), temporal (T7, T8), central (C3, Cz, C4), centroparietal (CP1, CP2, CP5, CP6), parietal (P3, P4, P7, P8, Pz), occipital (O1, O2, Oz) and aggregated the MSE values of these clusters over five time scales. This resulted in 32 (8 spatial clusters x 4 averaged scales) aggregated MSE measures, which were used as input features for the prediction models together with the measures derived from Shannon entropy, from Fuzzy entropy, and with the microstate measures. Performances of prediction models were calculated as the Pearson correlation between predicted and observed intelligence scores. To control for influences of age, sex, and the number of removed epochs, those variables were regressed out with linear regression from the input features (brain signal complexity measures) and from the prediction targets (RAPM scores). Complexity measures and RAPM scores were z-standardized after regressing out the confounds, respectively.

At first, we aimed to test how much variance in intelligence we can explain with this multimodal approach. Therefore, the model was built and applied on the whole sample (without any cross-validation). Second, we tested whether a model build in one subset of the sample (the training sample) could predict individual intelligence scores in a withheld part of the sample (the test sample). Therefore, we implemented 10-fold cross-validation, i.e., the sample was divided into ten subsamples (ensuring equal distributions of intelligence scores via stratified folds) and the intelligence-relevant features (as described above) were selected in nine subsamples only and then applied to predict the intelligence scores of the withheld tenth sample. This step was repeated ten times to generate predicted intelligence scores for all subjects which can then be compared to the observed scores. Significance of the cross-validated model was assessed with a non-parametric permutation test. More in detail, 100 models with varying stratified sample divisions were trained and tested using the observed intelligence scores. Then, the mean performance of these models was evaluated against the performances of 1,000 models trained and tested on permutated intelligence scores (null models). Performances above the 95% confidence interval (*α* < .05) of these null model performances were considered as significant.

Finally, we aimed to evaluate the generalizability of our model to a completely different cohort of subjects (see below). Here, we selected the features in the whole main sample and used this selection to predict the intelligence scores of the external replication sample (see below). Again, significant prediction of intelligence was evaluated with a permutation test. Specifically, the performance of the prediction of the observed intelligence scores in the replication sample was tested against the prediction of permutated intelligence scores (1,000 iterations). Again, performances above the 95% confidence interval (*α* < .05) of null model performances were considered significant.

### External Replication

For testing the robustness and generalizability of our findings all analyses were repeated in an independent sample of 60 right-handed students from the Goethe University Frankfurt who were recruited via local advertisement (placate, flyers) and completed the experiment for monetary compensation or student credits. Students with a Major or Minor study subject in Psychology were excluded. All participants were right-handed, had self-reported normal or corrected-to-normal visual acuity and no history of psychiatric or neurological diseases. The procedures were approved by the local ethics committee (# 2015-201) and informed written consent according to the Declaration of Helsinki was obtained from all participants. EEG recordings took place in a sound-shielded room, instructions were similar as in the main sample, and a total of five minutes resting-state data were acquired. One participant was excluded due to EEG acquisition failure, one due to a missing RAPM score and demographic data, and one participant due to excessive EEG artifacts, leaving a final sample of *N* = 57 subjects (16 male, 41 female) with age between 18 and 33 years (*M* = 23.51, *SD* = 3.61). Intelligence was assessed in group settings (10-12 participants) with Raven’s Advanced Progressive Matrices (RAPM, Raven and Court, 1998). RAPM scores ranged from 12 to 35 (*M* = 24.60, *SD* = 4.70; see Fig. 1B for details on RAPM’s distribution of the replication sample). EEG data were recorded with 64 active Ag/AgCl electrodes (arranged in an extended 10–20 layout) using actiChamp amplifier (Brain Products GmbH, Gilching, Germany). FCz was used as online reference, and AFz served as ground. The sampling rate was 1,000 Hz, impedance levels were kept below 10 kOm, and a low pass filter of 280 Hz was applied during acquisition (notch filter off). Two electrodes were placed below the left (SO1) and the right (SO2) eye to record ocular artifacts, and mastoid electrodes were placed behind both ears (M1, M2). Preprocessing was performed similar as in the main sample. Due to the higher number of electrodes, different to the main sample, FastICA with 30 components was applied on the data to identify artefact components of five subjects and to remove similar artifact components in all subjects. Note that after preprocessing the 64 scalp electrodes were reduced to the same electrodes as used in the main sample (28 scalp electrodes) to allow for direct comparisons. Statistical analyses were performed similar as in the main sample, but all partial correlations were controlled for sex (Pearson correlation between intelligence and sex: *r* = .00, *p* = .978) in addition to age (Pearson correlation between intelligence and age: *r* = −.21, *p* = .124) and the number of removed epochs (Pearson correlation between intelligence and the number of removed epochs: *r* = .03, *p* = .801). Note that for microstate analyses, the group microstates of the main sample were backfitted to the individual EEG-signals of the participants in the replication sample.

### Data and Code Accessibility

Analyses were conducted using Python 3.8. The code/software described in the paper is freely available online at https://github.com/jonasAthiele/BrainComplexity_Intelligence, https://doi.org/10.5281/zenodo.7258768. The code is available as Extended Data. The raw data can be accessed from the authors by reasonable request.

## Results

### Intrinsic Brain Signal Complexity

Group-average entropy measures as well as their standard deviation between participants are illustrated in Figure 3. Overall, mean and standard deviation of channel-wise Shannon entropy (Fig. 3A), channel-wise Fuzzy entropy (Fig. 3B) and channel-wise multiscale entropy (Fig. 3D) demonstrated obvious differences between different EEG-channels. Channel-wise means and standard deviations of Shannon entropy were descriptively higher than means and standard deviations of channel-wise Fuzzy entropy. Means of sample entropy increased between scale 1 and 5 and remained relatively stable from scale 5 to 20 (see multiscale entropy of GFP, Fig. 3C, and channel-wise multiscale entropy, Fig. 3D). The five group microstates extracted from the EEG data are illustrated in Figure 4A. Microstate patterns of resting-state EEG have been demonstrated to be highly reproducible between studies (Michel and Koenig, 2018) and can be categorized into established classes. The here extracted five microstates (Fig. 4A) map well onto the classes A, B, C, D, and F as established in previous studies (Custo et al., 2017; D’Croz-Baron et al., 2021; Férat et al., 2022). Means and standard deviations of microstate measures are shown in Figure 4B,C. Again, also these measures varied markedly between specific measures and between different microstates.

**Figure 3.**
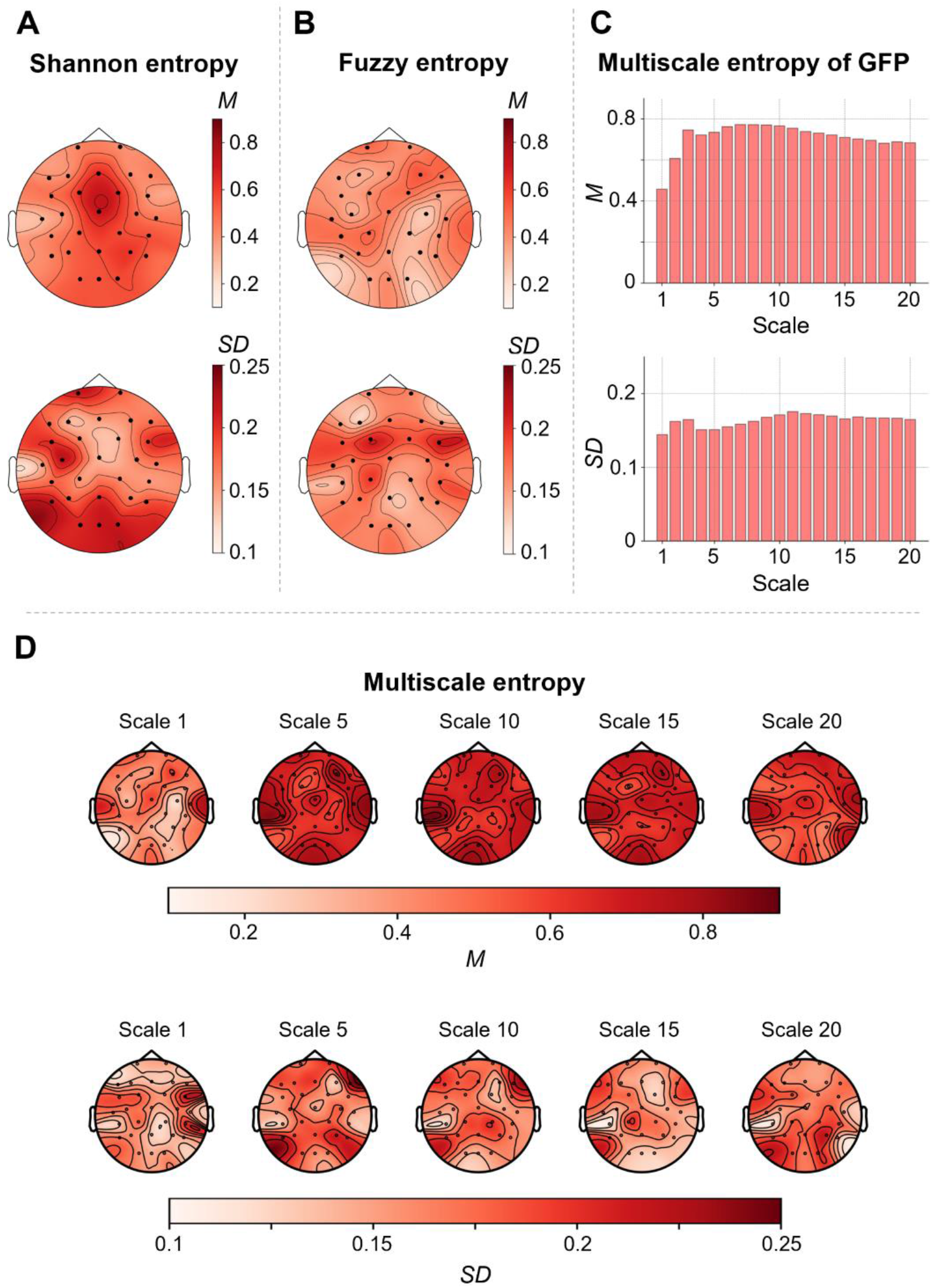
Entropy measures as indicators of brain signal complexity. Mean values (*M*) and standard deviations (*SD*) of three different entropy measures were calculated from the time series of neural activation as measured with resting-state electroencephalography (EEG) at 28 scalp electrodes. Mean and standard deviations of all three measures (normalized between 0 and 1) were calculated across 144 healthy adult participants. For illustration, values were interpolated and color-coded (see color bars). **A**, Mean and standard deviation of Shannon entropy computed on 28 scalp electrodes (represented as dots) separately. **B**, Mean and standard deviation of Fuzzy entropy of 28 scalp electrodes. **C**, Means and standard deviation of multiscale entropy (MSE), indexing the sample entropy at different coarse-grained time series (scales), computed on the global field power (GFP) of 28 scalp electrodes for the time scales 1 to 20. **D**, Mean and standard deviation of MSE of 28 scalp electrodes for different time scales (1,5,10,15, and 20). *Figure Contributions:* JT and KH designed research; JT, AR, KH performed research; JT & AR analyzed data.

**Figure 4.**
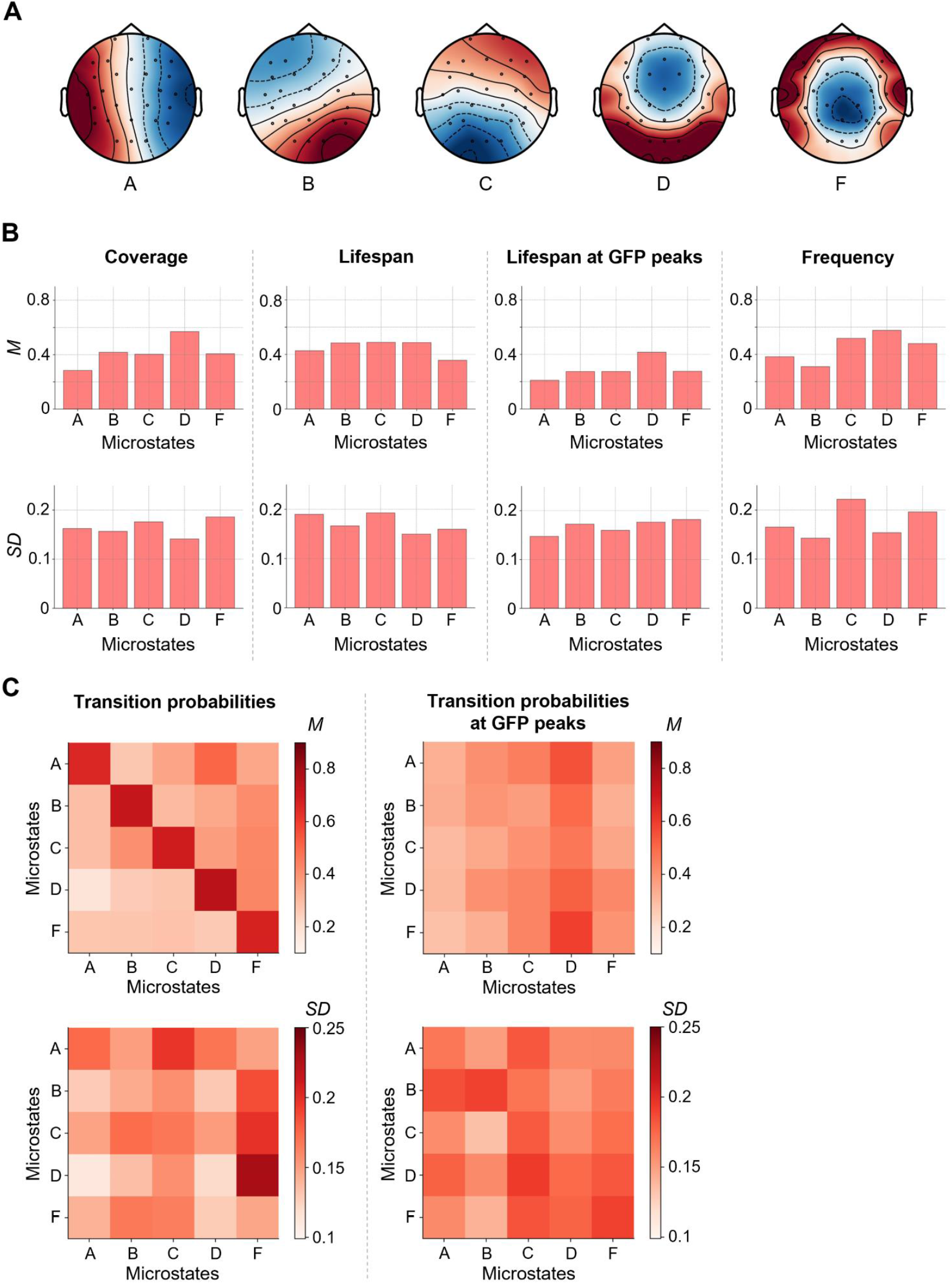
Microstate measures as indicators of brain signal complexity. Brain signals of 144 participants were clustered over time into five individual spatial mean maps (microstates). The individual mean maps of all participants were then clustered into five group microstates (**A**). These group-average microstates were then mapped back onto the individual electroencephalography (EEG) time series resulting in a sequence of group microstates for each individual. Based on these sequences, different measures were calculated (coverage, lifespan, lifespan at peaks of the global field power, frequency, and transition probabilities of microstates, see also Figure 2). For calculating means and standard deviations, all measures were normalized between 0 and 1. **B**, Means (top row) and standard deviations (bottom row) of individual coverage, lifespan, lifespan at GFP peaks, and frequency of group microstates. **C**, Maps of means and standard deviations of individual probabilities to stay in a specific microstate or to transition into another specific microstate calculated on the whole time-series (left) and on time points with GFP peaks only (right). Values are color-coded (see color bars). *M*, mean; *SD,* standard deviation; GFP, global field power. *Figure Contributions:* JT and KH designed research; JT, AR, KH performed research; JT & AR analyzed data.

An explorative factor analysis was conducted to examine the covariance structure of different brain signal complexity measures. Results are illustrated in Figure 5. As indicated by parallel analysis, the appropriate number of the factors to be extracted was 17 and the total amount of variance explained by these 17 factors was 87.12 % (Fig. 5A). Figure 5B lists all 17 extracted factors with the ten variables expressing the highest loadings onto these factors. For example, for the three factors explaining most variance, variables with highest loadings were MSE in coarser time scales (channels CP1, CP2, Fz; factor 1), MSE in finer time scales (channels FC1, FC2, FC5, Fz, CP5, CP6, C3, C4, P7; factor 2 and factor 3), and Fuzzy entropy (channels C3, FC5, FC6; factor 3).

**Figure 5.**
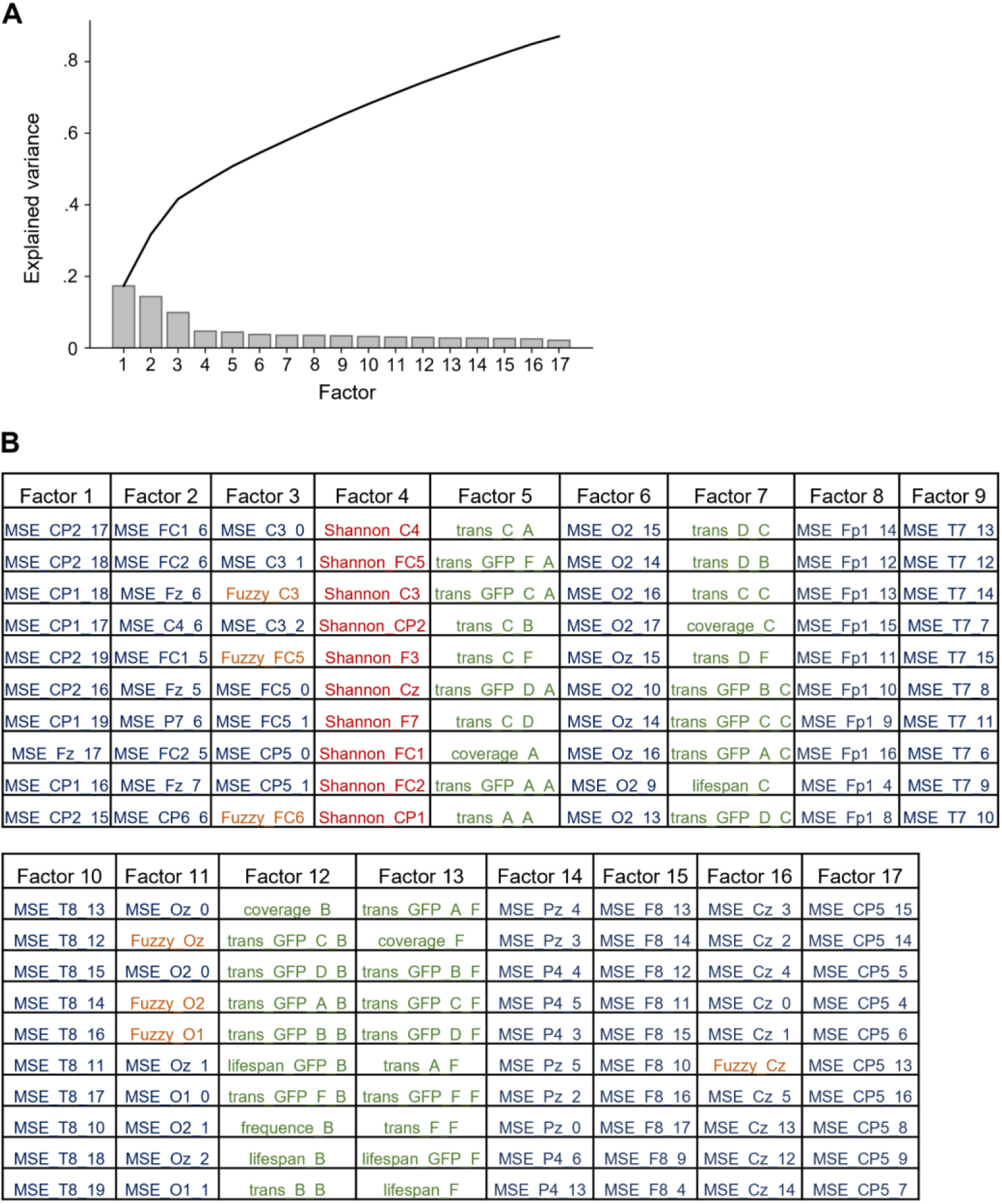
Measures of brain signal complexity can be grouped into 17 latent factors. Latent factors were extracted from all computed measures of brain signal complexity (see Methods) by exploratory factor analysis using oblique rotation and minimal residual minimization. To define the optimal number of factors, parallel analysis was used in accordance with Horn (1965). **A**, Proportion of total variance explained by each factor. The black curve shows the accumulation of explained variance. **B**, Extracted factors with (for illustration purposes) the ten brain signal complexity measures with highest loadings on the respective factor. Measures of multiscale entropy (MSE) are written in blue font, Shannon entropy measures in red, Fuzzy entropy measures in orange, and parameters derived from the microstate analysis are depicted in green. GFP, global field power. *Figure Contributions:* JT and KH designed research; JT, AR, KH performed research; JT & AR analyzed data.

### The Association Between Intelligence and Brain Signal Entropy Depends on Spatial and Temporal Scales

On a whole-brain level, we observed no significant associations between intelligence and Shannon or Fuzzy entropy (computed on the GFP signal; Shannon: *r* = −.09, *p* = .30; Fuzzy: *r* = −.05, *p* = .58). Channel-specific Shannon entropy values were mostly negatively associated with intelligence, but no association reached statistical significance (Fig. 6A). Fuzzy entropy showed a similar pattern, with some channel-specific associations reaching statistical significance when not correcting for multiple comparisons: Cz (*r* = −.21, *p* = .014, *p(adj)* = .129), CP1 (*r* = −.23, *p* = .007, *p(adj)* = .129), CP2 (*r* = −.21, *p* = .012, *p(adj)* = .129) and Pz (*r* = −.17, *p* = .045, *p(adj)* = .318, Fig. 6B). In the replication sample most associations were also negative, however, the specific channels showing strongest associations as well as the strengths of associations differed between the samples (see Fig. 6A,B).

**Figure 6.**
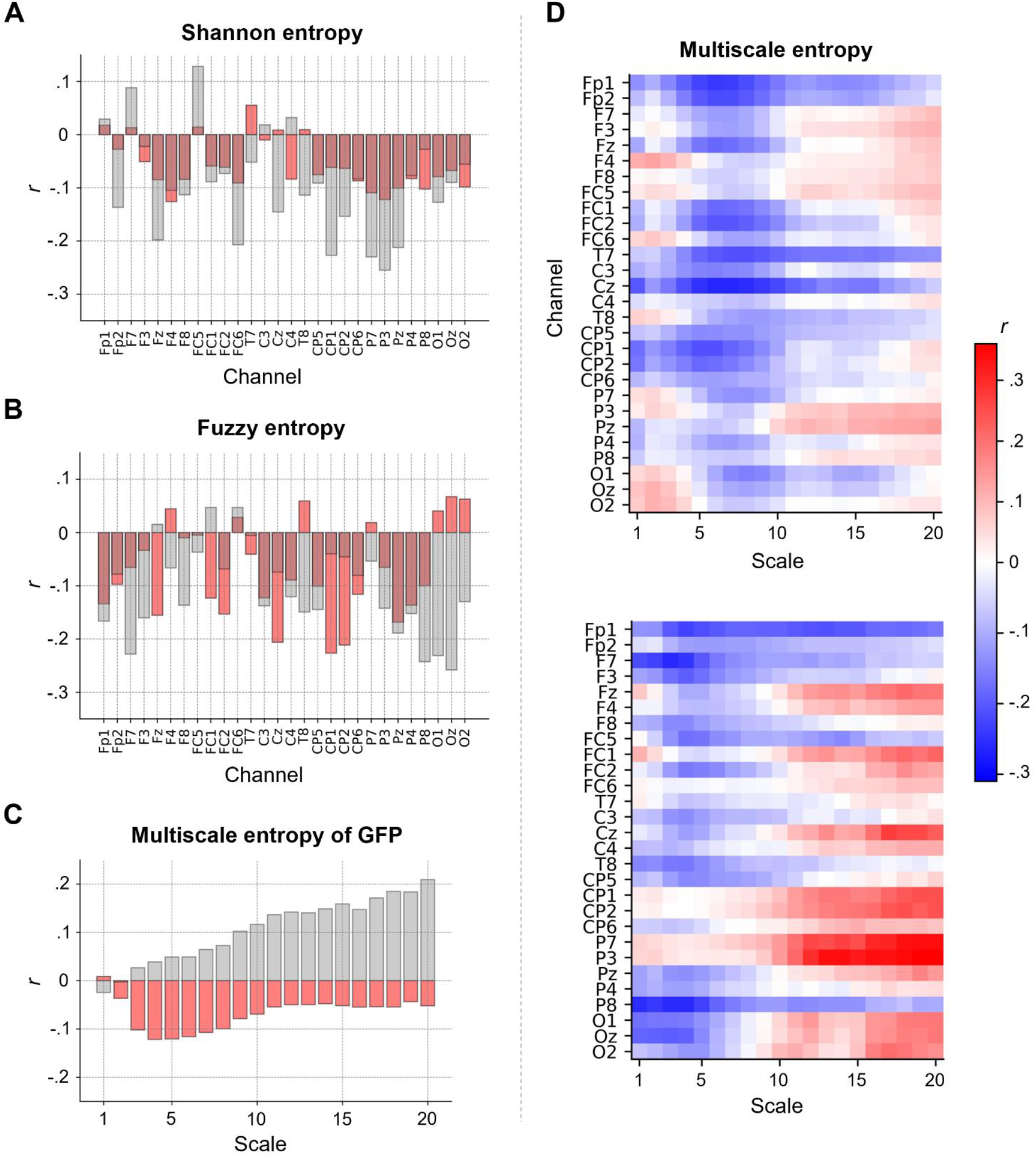
The association between intelligence and intrinsic brain signal entropy depends on electroencephalography (EEG) channel, time scale and study sample. Pearson correlations *r* (controlled for age, sex, and number of removed epochs) between intelligence (RAPM; Raven and Court, 1998) and (**A**) Shannon entropy for each EEG channel, (**B**) Fuzzy entropy for each EEG channel, and (**C**) multiscale entropy (MSE), indexing the sample entropy at different coarse-grained time series (temporal scales), of the global field power (GFP) at time scales 1 to 20. Associations found in the main sample (*N* = 144) are depicted in red, associations found in the replication sample (*N* = 57) are illustrated in gray. **D**, Pearson correlations *r* (controlled for age, sex, and number of removed epochs) between intelligence and MSE computed for the time scales 1 to 20 and each EEG channel. Upper panel: Main sample. Lower panel: Replication sample. Associations without confound corrections are shown in Extended Data Figure 6-1. *Figure Contributions:* JT and KH designed research; JT, AR, KH performed research; JT & AR analyzed data.

Whole-brain multiscale entropy (MSE, computed on the GFP signal) was on all but scale 1 negatively associated with intelligence in the main sample (without reaching statistical significance) while primarily positively associations were observed in the replication sample (see Fig. 6C). Channel specific MSE values were negatively associated with intelligence (only significant when uncorrected for multiple comparisons) for channels Fp1, Fp2, Fz, FC1, FC2, T7, Cz, CP1, and CP2, especially (but not exclusively) at finer time scales (scales 4-9), while we observed a non-significant trend towards positive associations for parietal (P3, Pz) and frontal channels (e.g., F3, FC5) at coarser time scales (scales 10-20; Fig. 6D). Similar tendencies were observed in the replication sample (Fig. 6D). However, while the patterns of associations between intelligence and sample entropy were significantly correlated between both samples (for all single scales and channels: *r* = .29, *p* < .001; for spatially and temporally related clusters that were used in the prediction models: *r* = .45, *p* = .001), associations of specific channels and time scales differed markedly. However, the overall pattern of lower entropy of fronto-central channels at finer time scales and tendencies towards higher entropy of fronto-parietal channels at coarser time scales to be associated with higher intelligence was observed in both samples. Associations between intelligence and entropy measures without controlling for confounds are illustrated in Extended Data Figure 6-1 and demonstrate that there were no substantial effects on the results by controlling for confounds.

### Intelligence is Associated with Two Specific EEG Microstates

Intelligence was positively correlated with the coverage (*r* = .20, *p* = .018, *p(adj)* = .077), lifespan (*r* = .20, *p* = .017, *p(adj)* = .077) and lifespan at GFP peaks (*r* = .20, *p* = .019, *p(adj)* = .077) of microstate A, the transition probability (all time points) from microstates A (*r* = .22, *p* = .010, *p(adj)* = .089), and C (*r* = .18, *p* = .036, *p(adj)* = .23) to microstate A (note that a transition probability from a specific microstate into the same microstate expresses the probability that the microstate does not change), as well as with the transition probability calculated at GFP peaks from microstate A (*r* = .22, *p* = .008, *p(adj)* = .048), B (*r* = .23, *p* = .007, *p(adj)* = .048), C (*r* = .22, *p* = .010, *p(adj)* = .048), and D (*r* = .20, *p* = .017, *p(adj)* = .061) to microstate A (Fig. 7). In contrast, intelligence was negatively correlated with the coverage (*r* = −.22, *p* = .009, *p(adj)* = .077) and frequency (*r* = −.21, *p* = .013, *p(adj)* = .077) of microstate C, the transition probabilities (all time points) from microstate A (*r* = −.21, *p* = .011, *p(adj)* = .089), and F (*r* = −.22, *p* = .010, *p(adj)* = .089) to microstate C as well as the transition probabilities (measured at GFP peaks only) from microstate A (*r* = −.20, *p* = .016, *p(adj)* = .061), D (*r* = −.24, *p* = .005, *p(adj)* = .048), and F (*r* = −.26, *p* = .002, *p(adj)* = .048) to microstate C (Fig. 7). All other measures as well as the number of GFP peaks did not show any significant correlations (all *p* > .05). In the replication sample, intelligence was associated in the same direction as in the main sample with the coverage, lifespan, and lifespan at GFP peaks of microstate A, coverage and frequency of microstate C, transition probabilities (all time points) from microstate A, and C to microstate A and from microstate A, and F to microstate C as well as with the transition probability at GFP peaks from microstate A to microstate A and from microstate A, D, and F to microstate C. However, overall, the strengths of associations differed between samples and few associations were also of opposing direction (Fig. 7). Associations between intelligence and microstate measures without controlling for confounds are illustrated in Extended Data Figure 7-1 and reveal that there were no substantial effects on the results by controlling for confounds.

**Figure 7.**
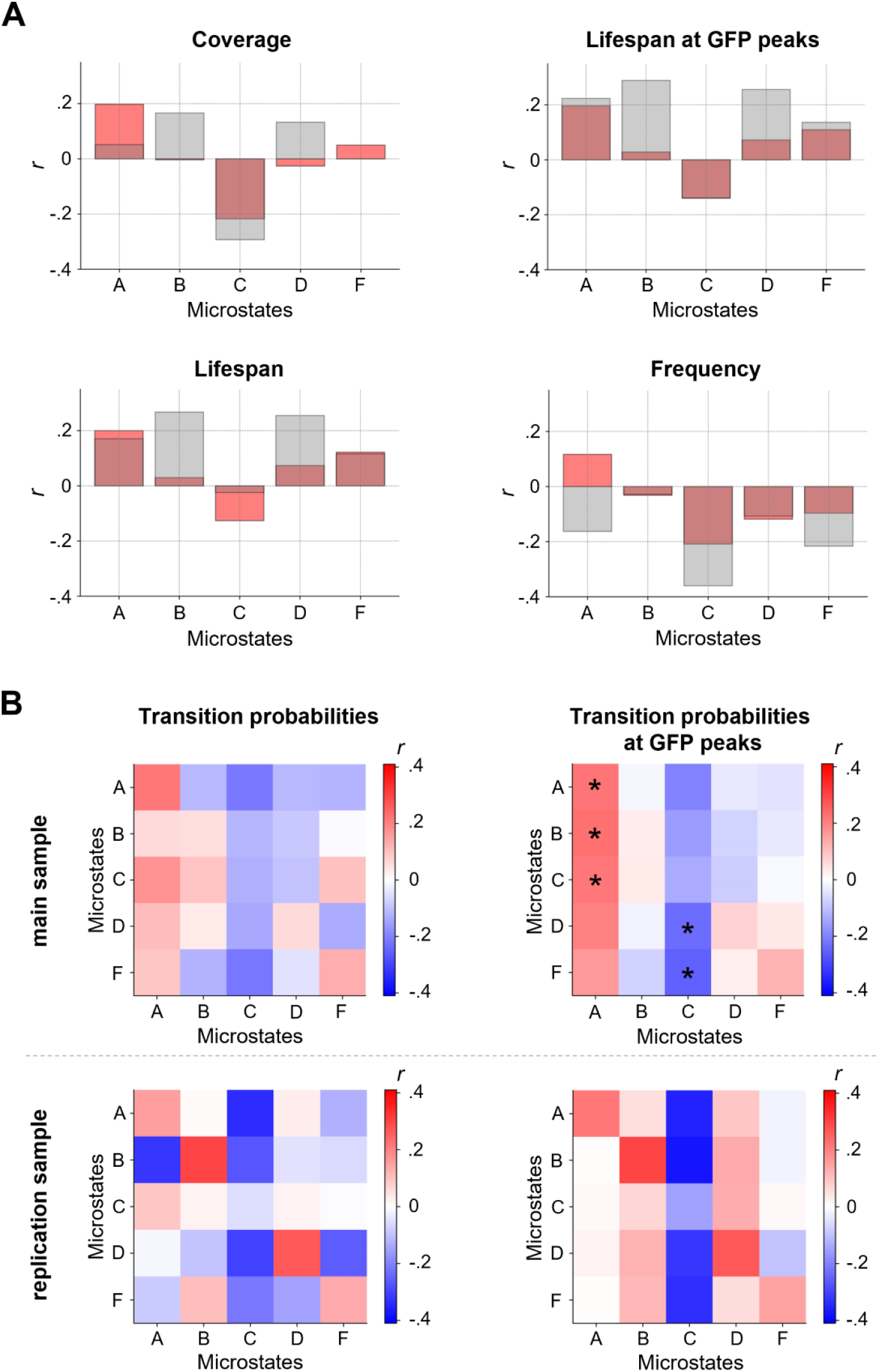
Intelligence is associated with two specific electroencephalography (EEG) microstates. Pearson correlations *r* between intelligence and characteristics of group microstates. Individual brain signals of 144 participants were clustered over time into five individual spatial mean maps (microstates). The individual mean maps of all participants were then clustered into five group microstates (see Fig. 4A). **A**, Pearson correlations (controlled for age, sex, and number of removed epochs) between individual intelligence scores (RAPM; Raven and Court, 1998) and variations in coverage, lifespan, lifespan at GFP peaks, and frequency of group microstates when mapped back onto the individual time series of neural activation from participants of the main sample (red, *N* = 144) and the replication sample (gray, *N* = 57). **B**, Pearson correlations (controlled for age, sex, and number of removed epochs) between intelligence (RAPM, Raven and Court, 1998) and individual-specific transition probabilities between group microstates (probabilities to stay in a specific microstate or transitioning into another specific microstate) when mapped back onto individuals’ time series from the main sample (top row) and the replication sample (bottom row), for transition probabilities on the whole time series (left) and on time points with GFP peaks only (right). FDR corrected p-values < .05 are marked with asterisks. GFP, global field power. Associations without confound corrections are shown in Extended Data Figure 7-1. *Figure Contributions:* JT and KH designed research; JT, AR, KH performed research; JT & AR analyzed data.

### Predicting Intelligence from Multimodal Brain Signal Complexity

To test if a combination of multiple measures of brain signal complexity can amplify the observed associations with intelligence, we firstly constructed a multimodal model to explain the variance in intelligence scores within the complete main sample. The combined average values of the variables positively and negatively correlated with intelligence (*p* < .05, uncorrected) served as model features (see Methods). Positively correlated variables were: coverage, lifespan and lifespan at GFP peaks of microstate A, transition probabilities from microstates A, and C to microstate A, and transition probability at GFP peaks from microstates A, B, C, and D, to microstate A. Negatively correlated variables were: frontopolar MSE scale 6-10, temporal MSE scale 6-10, central MSE scale 6-10, Fuzzy entropy of channel Cz, CP1, CP2, and Pz, coverage and frequency of microstate C, transition probabilities from microstates A, and F to microstate C, as well as the transition probabilities at GFP peaks from microstates A, D, and F to microstate C. This within-sample model (without any cross-validation) reached statistical significance (*r* = .31, *p* < .001, see Fig. 8A) indicating that a significant portion of variance in intelligence scores can be explained by a combination of multimodal brain complexity features.

**Figure 8.**
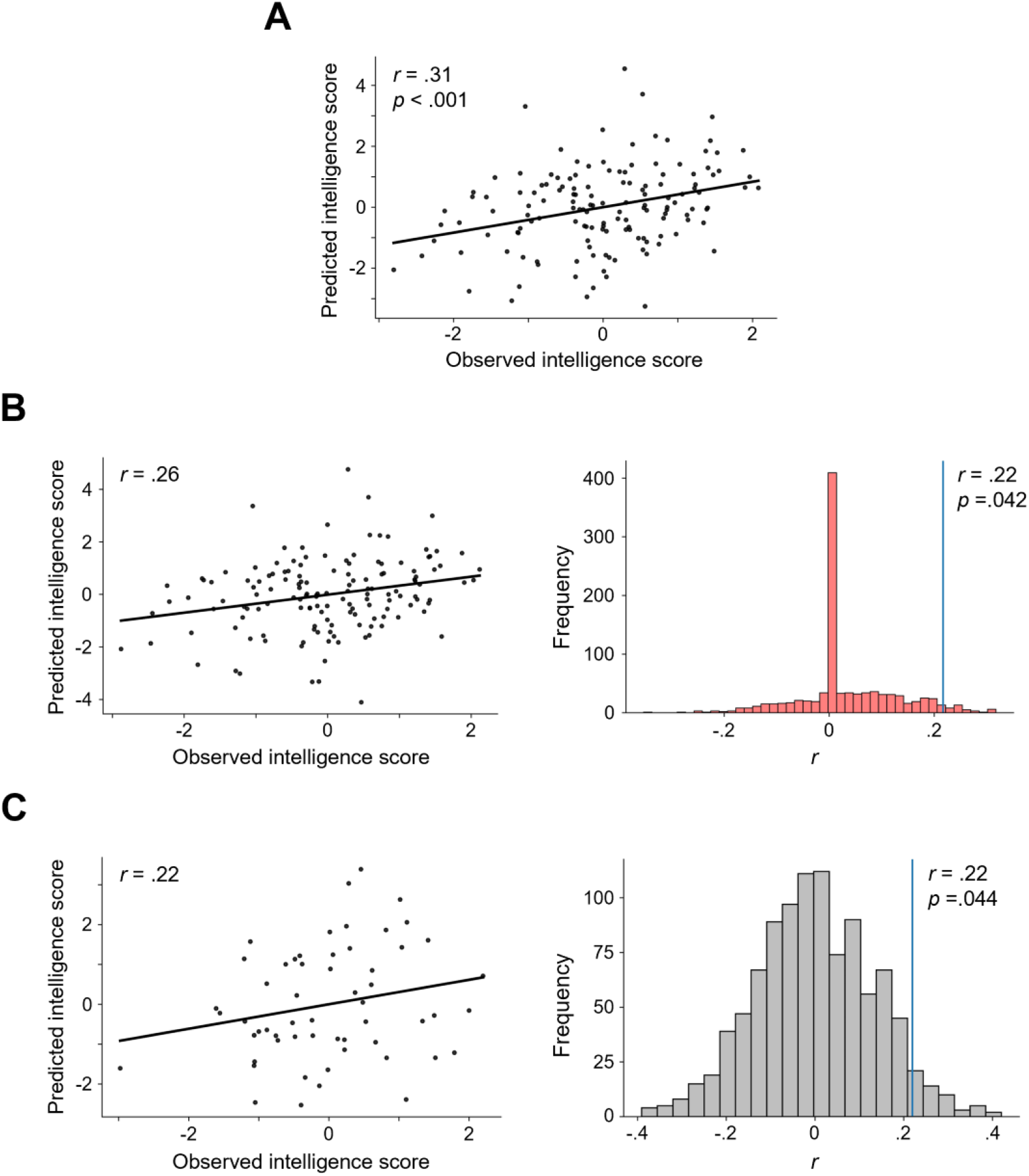
Multimodal brain signal complexity predicts individual intelligence (RAPM, Raven and Court, 1998) scores. Model performances were assessed via Pearson correlation *r* between the observed and predicted intelligence scores. Standardized residuals (controlled for age, sex, and number of removed epochs) of intelligence scores and complexity measures were used. **A**, Explanation: Model to explain variation in individual intelligence scores in the main sample (*N* = 144) from measures of brain signal complexity positively or negatively correlated (*p* < .05, uncorrected) with intelligence. **B**, In-sample prediction: Left: Observed vs. predicted intelligence scores based on 10-fold internal cross-validation within the main sample. Predicted intelligence scores result from a model based on measures of brain signal complexity positively and negatively correlated (*p* < .05, uncorrected) with intelligence. Results of the model with the highest accuracy (from 100 different stratified sample divisions). Right: Result of a permutation test for testing significance of the prediction. The correlation of the prediction model (blue vertical line) was computed as average of correlations between predicted and observed intelligence scores from 100 models (with different stratified sample divisions) using internal 10-fold cross-validation. This average correlation was then tested against model performances of models constructed on the basis of permutated intelligence scores (1000 times, null models, histogram). Note that the high frequency of zero correlations occurred as a correlation of zero was automatically set if no measure of brain signal complexity correlated significantly with the permutated RAPM scores (*p* < .05, uncorrected). **C**, Out-ofsample prediction: Left: Observed vs. predicted intelligence scores in the replication sample (*N* = 57). Predicted intelligence scores for the replication sample resulting from a model that was constructed on the main sample, i.e., model in (**A**). Right: Permutation test for testing the significance of this prediction. The true model performance (blue vertical line) was tested against predictions of the same model for 1000 permutated intelligence scores within the replication sample. *Figure Contributions:* JT and KH designed research; JT, AR, KH performed research; JT & AR analyzed data.

Secondly, we tested whether individual intelligence scores can not only be explained but also be predicted by our model. Therefore, we selected features on only one part of the sample and applied this model to the withheld part of the sample (internal 10-fold cross-validation). The model could significantly predict individual intelligence scores (correlation between predicted and observed intelligence scores: *r* = .22; *p* = .042 by permutation test, see Fig. 8B).

Finally, we tested whether a model trained on data from the main sample can also predict individual intelligence scores in a completely independent sample (out-of-sample prediction). This was indeed the case. Predicted and observed intelligence scores of the replication sample (external test sample) correlated significantly with *r* = .22, *p* = .044 (by permutation test), overall suggesting high generalizability of this multimodal approach (see Fig. 8C).

### Post-hoc Analyses

To gain additional exploratory insights into the associations between intelligence and intrinsic brain signal complexity beyond those provided by established and frequently used metrics, we created three additional measures capturing complementary aspects of brain signal complexity and evaluated their relationship with intelligence. Specifically, brain signal variety was estimated by the variety of individual microstates and the proportion of explained variance of the EEG signals by microstates. Firstly, we compared all subject-specific microstates with each other and tested whether their spatial similarity was related to intelligence. Spatial similarity was operationalized as the average of Pearson correlations between the vectorized spatial maps of each of two microstates. We observed a significant negative association of *r* = −.20, *p* = .019 suggesting less similarity (higher variability) between individual microstates in participants with higher intelligence scores. Secondly, we computed the total amount of variance in the subject-specific EEG signal that can be explained from the five extracted subject-specific microstates and tested whether this amount is associated with individual differences in intelligence. The average (across participants) total amount of explained variance was *M* = 63% (*SD* = 6%, range = 44% – 83%). However, this amount was not significantly correlated with intelligence (*r* = −.15, *p* = .071). Finally, we assessed the total amount of variance in the subject-specific EEG signal that can be explained by the five group microstates. Across participants the total amount of variance explained was *M* = 56% (*SD* = 10%, range = 22% – 76%) and, interestingly, we observed that in people with higher intelligence scores less variance could be explained by the group microstates (*r* = −.21, *p* = .011).

## Discussion

We showed that multiple measures capturing the complexity of intrinsic brain dynamics are associated with human intelligence. Specifically, Shannon, Fuzzy, and multiscale entropy of resting-state EEG were compared with features of established microstates and factor-analytical results point towards the existence of overlapping as well as separate information captured by the different measures. Further analyses revealed that associations between brain signal complexity and intelligence not only vary between different measures but do also critically depend on the considered EEG channel and on the focused temporal scale, thus only little variance in intelligence may be explained by unimodal approaches. Therefore, we finally combined different measures into a multimodal model considering different measures, channels, and time scales simultaneously. This model allowed significant prediction of individual intelligence scores in the main sample as well as in a completely independent sample.

At first, our study reveals that the complexity measures calculated for different EEG channels and on different temporal scales, can be grouped into 17 latent factors. These factors cluster in respect to spatial information and time scale as well as to the kind of measure. For instance, Fuzzy entropy and MSE measures were clearly differentiated from Shannon entropy and microstate characteristics, respectively. These observations demonstrate that different amounts of variance in the complexity of the resting-state EEG signal are captured by different measures, and suggest that physiologic signal complexity might be too manifold to be captured in its entirety by only one single metric (Goldberger et al., 2002). Conclusively, also for investigating associations between brain signal complexity and human traits, it might be more appropriate to use multiple complexity measures.

When computed on the global field power, no entropy measure was significantly associated with intelligence. Channel-specific Shannon and Fuzzy entropy were descriptively negatively associated with intelligence but did not reach statistical significance. Sample entropy of fronto-central channels demonstrated non-significant negative associations with intelligence at finer timescales, while sample entropy of fronto-parietal channels at coarser time scales showed non-significant tendencies towards positive associations. Together, the finding of lower entropy at finer time scales and tendencies towards higher entropy at coarser time scales to be associated with higher intelligence is in line with the findings of Dreszer et al. (2020) suggesting differences in local (linked to entropy at finer time scales) versus global (as reflected by entropy at coarser time scales) aspects (Courtiol et al., 2016; McIntosh et al., 2014; Vakorin et al., 2011) of intelligence-related information processes. Interestingly, it has been shown that entropy at finer time scales increases while entropy at coarser time scales decreases with increasing age (McIntosh et al., 2014), which is in regard to our observations, plausible as fluid intelligence decreases with increasing age (Ghisletta et al., 2012; Salthouse, 2010; Schaie, 1994). We, therefore, would like to encourage future studies to test whether both associations exist rather independent of each other or whether intrinsic brain signal entropy mediates the association between increasing age and reductions in intelligence.

The analysis of EEG microstates revealed intelligence-related differences in respect to two specific microstates. Specifically, higher intelligence scores were associated with less dominance of microstate C as indicated by significantly less transitions into microstate C and descriptive trends towards lower coverage and frequency - observations which replicated earlier reports (Liu et al., 2020; Santarnecchi et al., 2017). As microstate C is suggested to reflect increased activity of the default-mode (or task-negative) network (DMN, Michel and Koenig, 2018), and effective suppression of DMN activity is proposed to be essential for proper cognitive functioning (Anticevic et al., 2012; Sidlauskaite et al., 2016; Sonuga-Barke and Castellanos, 2007), this observation may indirectly point towards an intrinsic advantage of higher intelligent people for more effective DMN suppression. Further, higher intelligence was associated with higher presence of microstate A, reflected by more transitions into microstate A and trends towards higher coverage and lifespan. Microstate A has been related to reduced activity of the temporal network (Michel and Koenig, 2018) and especially to areas implicated in phonological processing (Britz et al., 2010), which is consistent with the observation that Microstate A is more present during visualization tasks expectedly implying inhibition of left-hemispheric language areas (Milz et al., 2016). However, implications of microstate patterns on cognition are far away from being completely understood and more research is needed to clarify whether and to which extend a higher presence of microstate A may possibly reflect intrinsic dispositions for verbal processing (Dreszer et al., 2020) or visualization (Milz et al., 2016).

Importantly, according to the criteria of Cohen (1988) all observed effect sizes (*r* ~ .2) can be considered as small and only few out of all calculated measures reached statistical significance, when correcting for multiple comparisons. These results contribute to the current debate about the effect size to be expected in investigations on brain-behavior relations (Marek et al., 2022; Rosenberg and Finn, 2022) in demonstrating that the combination of crossvalidation (Cwiek et al., 2022; Sui et al., 2020) and multimodal analyses approaches can identify robust brain-behavior relations despite sample sizes that lie clearly below thousand. We compared three forms of analyses (explanation of intelligence scores, internal crossvalidation, and out-of-sample prediction: prediction of intelligence scores in a replication sample with the model constructed in the main sample) to show, in line with Cwiek et al. (2022), that cross-validation reduces the overall effect size markedly as compared to explanation (*r* = .31 to *r* = .22). Notably, that internally cross-validated effect sizes reflect more realistic estimates of the ‘true’ effect sizes (Yarkoni and Westfall, 2017) is supported by our out-of-sample-prediction in the independent sample (*r* = .22).

In both cases (internal cross-validation and out-of-sample prediction) our models could significantly predict individual intelligence scores. Features contributing to this model were frontopolar, central, and temporal sample entropy at finer timescales, centro-parietal Fuzzy entropy, and measures of microstate A and C, which again supports the relevance of these aspects for intelligence. The involvement of multiple brain areas in associations between intelligence and brain signal complexity supports theories proposing a distributed network of brain regions associated with diverse cognitive functions as relevant for the explanation of individual differences in intelligence (Basten et al., 2015; Duncan, 2010; Jung and Haier, 2007).

Finally, post-hoc analyses revealed that higher intelligence scores were associated with higher variability in individual-specific microstates and with less variance explained by group microstates. Although, this could be interpreted as pointing towards a more diverse intrinsic brain network configuration in more intelligent people at time scales in the range of milliseconds, it is difficult to relate those findings to recent neuroimaging studies suggesting higher stability (less variability) of intrinsic brain activity in more intelligent people at much coarser time scales (Hilger et al., 2020). A pressing goal for future research is therefore to link those lines of research and to systematically assess how intelligence-related spatial configurations (microstates) of resting-state EEG relate to intelligence-related brain network reconfigurations as observed in fMRI studies (e.g., Thiele et al., 2022).

Several limitations and methodological aspects require consideration. At first, we compared a very high number of parameters describing different aspects of intrinsic brain signal complexity, making it difficult to control for multiple comparisons. Although this was necessary due to the exploratory hypothesis-generating purpose of our study, and we attempted to control for multiple comparisons appropriately, this may have reduced our ability to detect significant associations. Second, the age range of our sample was restricted (18 to 35 years). As spontaneous brain dynamics can be influenced by age (Goldberger et al., 2002), future studies should test whether and to which extend our findings may generalize to different age cohorts. Third, our study was restricted to a specific collection of complexity measures most established in this field of research. Evaluating interplays of additional (and more diverse) complexity measures may add further insights into intrinsic brain dynamics underlying intelligence. Finally, as observed associations between complex human traits and neural parameters were relatively small, we recommend future studies aiming to investigate associations between complex human traits and complex neural parameters a) to use large samples (*N* > 200) for capturing enough phenotypic variation and reach sufficed statistical power (> .95) to detect expectedly small effects (*r* ~ .20), b) to combine multiple neural parameters in multimodal models, c) to apply internal cross-validation to obtain realistic estimates of the generalization error, and d) to actually test the generalizability in a completely independent sample.

In sum, our study reveals that individual differences in a person’s cognitive ability level are reflected in the complexity of temporally highly resolved intrinsic brain dynamics. We demonstrated that different complexity measures capture separate but also overlapping information and that associations with intelligence vary across temporal and spatial scales. Finally, we showed that combining different measures into a single multivariate model allows to significantly predict individual intelligence scores from only five minutes of resting-state EEG data. Overall, our study highlights the potential of combining multimodal analysis approaches with internal cross-validation and out-of-sample prediction to reliably investigate how intrinsic brain dynamics might contribute to complex human traits.

## Acknowledgements

The authors thank Ines Jetzinger, Amelie Schirmer, and Kilian Rolle for contributions in EEG data collection, Christian Fiebach (Frankfurt University) for providing resources for data acquisition of the replication sample, Jona Sassenhagen for help in data acquisition of the replication sample, Maren Wehrheim for comments on the manuscript, and all study participants for their participation.

## Conflict of interest

Authors report no conflict of interest.

## Funding sources

This work was supported by the German Research Foundation (DFG, grant number HI 2185-1/1 to K.H.), and the Heinrich-Böll Foundation (funds from the Federal Ministry of Education and Research, grant number P145957 to J.T.).

---

**Extended Data Figure 6-1.**
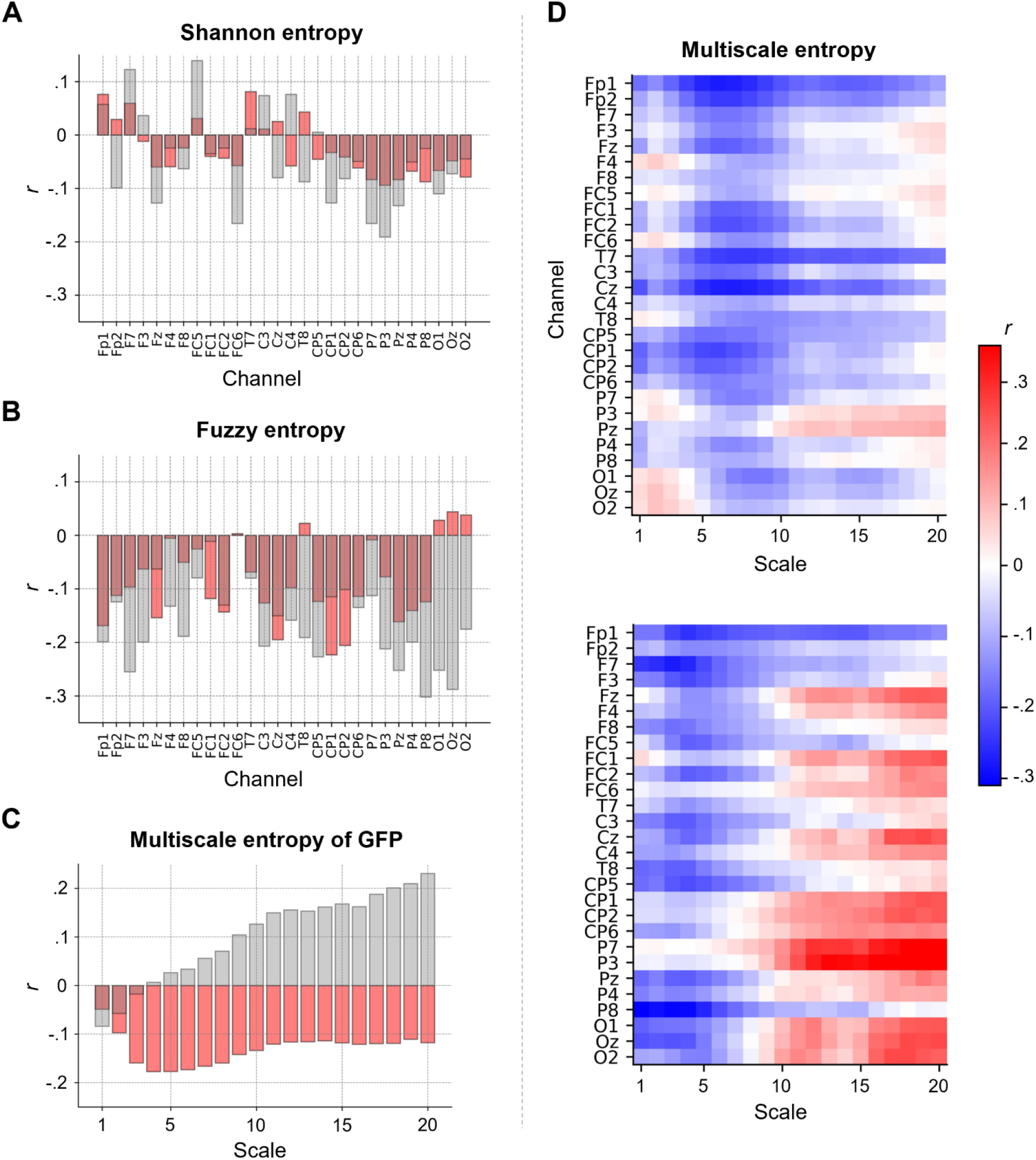
Scripts for the analyses used in the paper. The association between intelligence and intrinsic brain signal entropy depends on electroencephalography (EEG) channel, time scale and study sample. Pearson correlations *r* (not controlled for age, sex, and number of removed epochs) between intelligence (RAPM; Raven and Court, 1998) and (**A**) Shannon entropy for each EEG channel, (**B**) Fuzzy entropy for each EEG channel, and (**C**) multiscale entropy (MSE), indexing the sample entropy at different coarse-grained time series (temporal scales), of the global field power (GFP) at time scales 1 to 20. Associations found in the main sample (*N* = 144) are depicted in red, associations found in the replication sample (*N* = 57) are illustrated in gray. **D**, Pearson correlations *r* (not controlled for age, sex, and number of removed epochs) between intelligence and MSE computed for the time scales 1 to 20 and each EEG channel. Upper panel: Main sample. Lower panel: Replication sample. *Figure Contributions:* JT and KH designed research; JT, AR, KH performed research; JT & AR analyzed data.

**Extended Data Figure 7-1.**
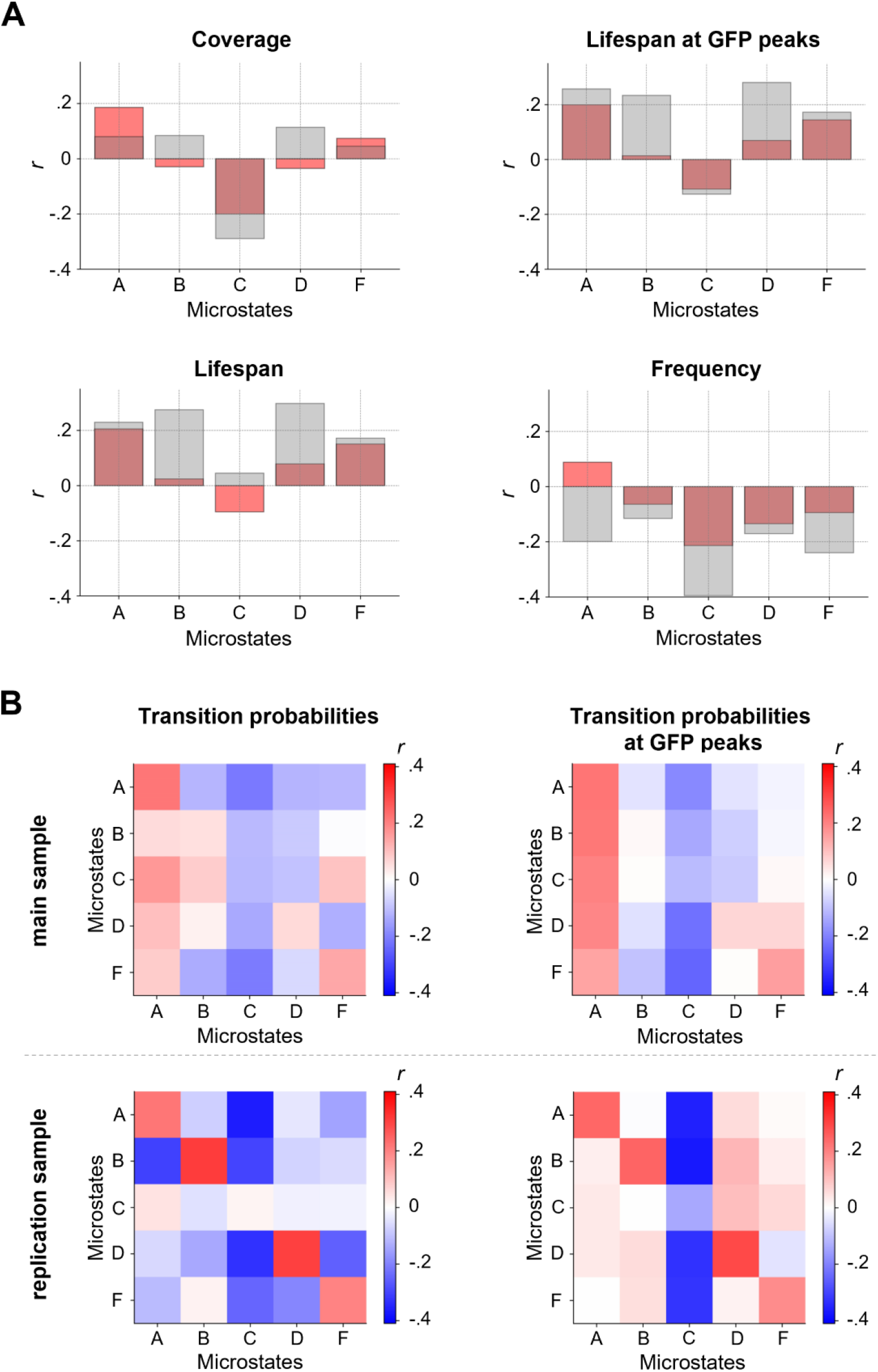
Intelligence is associated with two specific electroencephalography (EEG) microstates. Pearson correlations *r* between intelligence and characteristics of group microstates. Individual brain signals of 144 participants were clustered over time into five individual spatial mean maps (microstates). The individual mean maps of all participants were then clustered into five group microstates (see Fig. 4A). **A**, Pearson correlations (not controlled for age, sex, and number of removed epochs) between individual intelligence scores (RAPM; Raven and Court, 1998) and variations in coverage, lifespan, lifespan at GFP peaks, and frequency of group microstates when mapped back onto the individual time series of neural activation from participants of the main sample (red, *N* = 144) and the replication sample (gray, *N* = 57). **B**, Pearson correlations (not controlled for age, sex, and number of removed epochs) between intelligence (RAPM, Raven and Court, 1998) and individual-specific transition probabilities between group microstates (probabilities to stay in a specific microstate or transitioning into another specific microstate) when mapped back onto individuals’ time series from the main sample (top row) and the replication sample (bottom row), for transition probabilities on the whole time series (left) and on time points with GFP peaks only (right). FDR corrected p-values < .05 are marked with asterisks. GFP, global field power. *Figure Contributions:* JT and KH designed research; JT, AR, KH performed research; JT & AR analyzed data.

## Notes

https://github.com/jonasAthiele/BrainComplexity_Intelligence

https://doi.org/10.5281/zenodo.7258768

### Competing Interest Statement

The authors have declared no competing interest.

### Summary of Updates

Our main changes include: - Clarifications on EEG preprocessing steps and on the explanation of entropy measures - A discussion on the similarity of intelligence-entropy associations between the main and replication sample - Additional control analyses to rule out that our results are biased by controlling for confounds

